# Integrin β1 optimizes diabetogenic T cell migration and function in the pancreas

**DOI:** 10.1101/277418

**Authors:** Gabriel Espinosa-Carrasco, Cécile Le Saout, Pierre Fontanaud, Aurélien Michau, Patrice Mollard, Javier Hernandez, Marie Schaeffer

**Affiliations:** INSERM U1183, Institute for Regenerative Medicine and Biotherapy, University of Montpellier, Montpellier, France; Institute of Functional Genomics, CNRS, INSERM U1191, University of Montpellier, F-34094 Montpellier, France

**Author notes:** These authors contributed equally to this work (Gabriel Espinosa-Carrasco, Cécile Le Saout). Co-corresponding authors, Primary corresponding author: Dr. Marie Schaeffer, Inserm U1191, Institute of Functional, Genomics, Montpellier 34094, France, Corresponding author: Dr. Javier Hernandez, Inserm U1183, Institute for Regenerative Medicine and Biotherapy, Montpellier 34295, France. Current address: Memorial Sloan Kettering Cancer Center Sloan Kettering Institute, Zuckerman Research Center, New York, NY 10021, USA.

**Keywords:** autoimmunity, T cell migration, Type 1 diabetes, imaging, *in vivo*

## Abstract

T cell search behavior is dictated by their need to encounter their specific antigen to eliminate target cells. However, mechanisms controlling effector T cell motility are highly tissue-dependent. Specifically, how diabetogenic T cells encounter their target beta cells in dispersed islets throughout the pancreas during autoimmune diabetes remains unclear. Using intra-vital 2-photon microscopy in a mouse model of diabetes, we found that CXCR3 chemokine downregulated CD8+ T cell motility specifically within islets, promoting effector cell confinement to their target sites. In contrast, T cell velocity and directionality in the exocrine tissue were enhanced along blood vessels and extra-cellular matrix fibers. This guided migration implicated integrin-dependent interactions, since integrin blockade impaired exocrine T cell motility. In addition, integrin β1 blockade decreased CD4+ T cell effector phenotype specifically in the pancreas. Thus, we unveil an important role for integrins in the pancreas during autoimmune diabetes that may have important implications for the design of new therapies.

## INTRODUCTION

Immune responses implicate sequential encounters between T cells and their specific (or cognate) antigen in different body compartments to ensure efficient T cell priming, activation, and antigen clearance (1,2). However, the frequency of naïve T cells specific for a given antigen is low (3), and antigen abundance in target tissues may be variable and/or spatially restricted. Thus, T cell search behavior is driven by the need to actively explore the environment and locate cognate antigens. Since T cell migration patterns depend on cell-intrinsic parameters, context-dependent micro-environmental chemotactic cues and tissue-dependent structural features (4,5), empirical studies are required to identify T cell search mechanisms in specific disease settings. Given the importance of T cell search strategies in target cell clearance (1,2), mechanisms involved may constitute promising new therapeutic targets.

Dynamics and mechanisms of T cell migration leading to initial antigen encounter in secondary lymphoid organs are best characterized (4,6,7). In lymph nodes (LNs), the frequency of naïve antigen-specific T cells is low (3) and migration patterns must optimize the likelihood of a productive encounter with a cognate antigen-bearing antigen presenting cell (APC). Hence, naïve T cells typically display a high velocity dependent on chemokines and interactions with dendritic cells (DCs) (8,9). They migrate following a “Brownian” random walk intrinsically encoded (7,10) and guided by a network of fibroblast reticular cells (FRCs) (11). This ensures efficient sampling of a multitude of APCs (6) to promote rare cognate antigen encounter and naïve T cells activation. Activated effector T cells with reprogrammed expression of adhesion molecules and chemokine receptors then migrate to peripheral tissues (12), where they usually accumulate in large numbers and need to search for their spatially-restricted cognate antigen (3), to maintain effector functions (13) and eliminate target cells (14).

While the unique LN architecture facilitates antigen-T cell encounters, peripheral tissue geometry and composition greatly impact T cell migratory patterns and speed (1,10,15–17). For instance, vascular network, APC networks, and the extra-cellular matrix (ECM) architecture influence T cell interstitial trafficking through physical or/and adhesive guidance (15,17–19). While adhesion-dependent mechanisms are not required for interstitial migration and T cell motility in LNs is integrin-independent (20,21), T cells are able to switch migration modes *in vitro* (22) and inflammation-mediated changes in ECM composition in peripheral tissues are able to induce integrin-dependent T cell trafficking (1). Thus, predicting disease-dependent mechanisms controlling T cell motility in the periphery remains impossible, although these may play a crucial role in target cell clearance (1,2).

During type 1 diabetes (T1D), an autoimmune disease leading to destruction of insulin-producing pancreatic beta cells, beta cell-specific T cells become activated in the draining lymph nodes (23). Effector T cells then migrate to the pancreas and extravasate both within islets (24) and at post-capillary venules in the exocrine tissue (14). Furthermore, effector T cells have been shown to displace from one islet to another (14). These observations indicate that migration of T cells in the exocrine tissue to reach dispersed target islets is essential for disease progression. However, mechanisms governing their motility remain unclear. Recent work in a viral induced mouse model of diabetes described diabetogenic T cell motility as a Brownian-type random walk around islets (14), whereas in NOD mice they appear to migrate along blood vessels (19). Given the extensive ECM remodeling and the key role of ECM organization in T1D pathogenesis (25), we sought to investigate mechanisms of effector T cell interstitial migration in the pancreas during T1D onset, using intravital 2-photon imaging in a mouse model of autoimmune diabetes.

## MATERIAL AND METHODS

### Ethical Statement

Animal studies were conducted according to the European guidelines for animal welfare (2010/63/EU). Protocols were approved by the Institutional Animal Care and Use Committee (CEEA-LR-1190 and −12163) and the French Ministry of Agriculture (APAFIS#3874).

### Mice

Mice were bred in specific-pathogen-free facility and housed in conventional facility during experimentation. The transgenic mouse model of diabetes (26,27) involved InsHA (28), Clone 4 TCR (MHC class I-restricted) (29), and HNT TCR (MHC class II-restricted) mice (30) (from Prof. Sherman, The Scripps Research Institute, San Diego, USA)(27), RIPmCherry mice (31) (from National Institute of Medical Research, London, UK), and β-actin-GFP and -CFP mice (Jackson Laboratory). Clone 4 TCR Thy1.1 x β-actin-GFP, HNT TCR Thy1.1 x β-actin-CFP, and InsHA x RIP-mCherry mice on BALB/c x C57BL/6 background 10-16 weeks old were used (27). Littermate males and females were used whenever possible and homogeneously mixed between experimental groups.

### T cell isolation, adoptive transfer and diabetes monitoring

Equal numbers (2-3×10^6^ cells/recipient) of naïve CD8^+^ and CD4^+^ T cells isolated from Clone 4 TCR Thy1.1 x β-actin-GFP and HNT TCR Thy1.1 x β-actin CFP mice, respectively, were injected i.v. into InsHA x RIPmCherry mice sub-lethally irradiated (4.5 Gy) 24 h before in a therapeutic irradiator (Varian), as described (27). Mice were used for intra-vital imaging, sacrificed at day 10 for T cell characterization or monitored for diabetes onset. Recipient mice blood glucose levels were measured using a glucometer (AccuCheck).

### *In vivo* antibody and peptide treatment

Anti-CXCR3 (armenian hamster IgG, BioXcell) or isotype control polyclonal armenian hamster IgG (BioXcell) were injected i.v. (300 µg/mouse) on day 8 after T cell transfer 1 h prior to imaging *in vivo*. Anti-β_1_ integrin (Hmβ1-1, eBioscience) or isotype control armenian hamster IgG H4/8 (eBioscience) (100 µg/mouse) were injected i.v. on day 8 after T cell transfer 1 h prior to imaging. GRGDS peptide or control reverse SDGRG peptide (Sigma) (500 µg/mouse) were injected i.v. 10 min prior to imaging. For characterization of donor T cells by FACS, anti-β1 integrin or isotype control antibody were injected i.p. on days 8 (200 µg/mouse) and 9 (100 µg/mouse) and mice killed at day 10 after T cell transfer.

### Surgery and intra-vital imaging

All experiments used normoglycemic mice. Animals were anesthetized by injection of ketamine/xylazine (0.1/0.02 mg/g). Pancreas was exteriorized by surgery as described (27,31). Fluorescence was visualized using a Zeiss 7MP 2-photon microscope adapted with an M Plan Apo NIR ×20 objective (0.4 NA, Mitutoyo). Excitation was achieved using a Ti:Sapphire Chameleon Laser (Coherent) tuned to either 820 nm (mCherry, mCherry-GFP-CFP excitation and second harmonic generation (SHG)), 850 nm (rhodamine-GFP-CFP) or 880 nm (GFP-CFP). Fluorescence was captured using GaAsP PMTs at 460-500 nm for CFP, 500-550 nm for GFP, 610-700 nm for mCherry and rhodamine, and < 410 nm for SHG. Surface islets (< 100 µm) were identified using mCherry or by light contrast. Tissue viability was verified by fluorescent dextran injection i.v. (27).

### Image data analysis

Stacks 150 to 250 µm thick (Z steps of 3 µm) were acquired every 30 s to 1 min during 10 to 27 min. Movies were stabilized using Huygens Essential (SVI). Measurements were performed in at least 3 independent experiments. Average velocities and mean squared displacements (MSD) of individual T cells were obtained using Imaris (Bitplane). Directionality indexes (ratio between the distance between start and end time points in a straight line and the total length of the migratory path) were calculated using a routine programmed in MATLAB (18). Similarly, T cell coordinates obtained using Imaris were imported in MATLAB to measure displacement of T cells towards or away from islet centroids, to project T cell orientation of displacement vectors on a circle, to calculate angle differences between T cells displacement vector projections on the XY plane and direction of vessels, and to generate graphs of XY projections of T cell tracks, using custom programs (available upon demand). Areas with similar infiltration (100-320 total number of tracks/0.05 mm^3^ imaging volume) were compared. Cell tracks lasting less than 4 min were excluded. No exclusion was made based on velocity.

To analyze migratory patterns of T cell populations, equations describing the major models of diffusion of particles (Y = B1*t +(B2*t)^α^ with α = 2 for directed or ballistic motion, α = 1 and B2 = 0 for Brownian random walk, 0 < α < 1 and B1 = 0 for sub-diffusive or anomalous random walk, 1 < α < 2 and B1 = 0 for Lévy-type super-diffusive random-walk, and Y = Plateau*(1-exp(-K*t) for confined motility) (32) applied to the description of T cells migration (4) were used as models of non-linear regression to fit mean squared displacement (MSD) increase over time in GraphPad Prism (t is time, B1 and B2 are fitting parameters, K is the constant rate). In each case, the model providing the best fit (highest R^2^) was chosen to describe the pattern of motility.

### Flow Cytometry

For T cell phenotyping, single cell suspensions from pancreatic lymph nodes or pancreas infiltrating cells were prepared and stained as described (26). For intracellular cytokine staining, T cells were restimuated *ex vivo* with HA-specific peptides during 5 h before staining as previously described (26). The mAbs used were: anti-CD61 (ITGβ3)-FITC, anti-CD51 (ITGαV)-PE, anti-CD49e (ITGα5)-APC, anti-CD183 (CXCR3)-Alexa Fluor 780, anti-CD29 (ITGβ1)-Pacific blue (BioLegend, San Diego, CA); anti-CD4-V500, anti-CD4-FITC, anti-CD90.1 (Thy1.1)-PerCP, anti-CD90.1 (Thy1.1)-V450, anti-CD8a-V450, anti-CD62L-APC, anti-IL-2-APC, anti-IFNΓ-PE (BD Pharmingen); anti-CD8a-APC-Alexa Fluor 780, anti-IL-17-Alexa Fluor, anti-CD25-APC-Alexa Fluor 780 and anti-KLRG1-PE-Cy7 (eBioscience). Cells were analyzed on a FACSCanto II or a LSR Fortessa apparatus using Diva software (BDB).

### Confocal imaging

Pancreas preparation and antibody labeling were as described (33). Antibodies used were: hamster anti-CD11c (clone N418 1:300, eBioscience); rat anti-F4/80 (clone MCA4976 1:200, BioRad), rabbit anti-insulin (1:500, Cell Signaling); rat anti-endomucin (1:500, Santa Cruz Biotechnology); rabbit anti-fibronectin (clone AB1942 1:5000, Chemicon); mouse anti-collagen I (1:300, Abcam). Nuclei were labeled using dapi (Sigma). One to four slices were randomly selected from > 3 animals/group. Images were acquired using a Zeiss LSM 780 confocal microscope and analyzed using Imaris (Bitplane) and ImageJ (NIH).

### Statistical analysis

Values are represented as mean ± SEM. Statistical tests were performed using GraphPad Prism. Normality was tested using D’Agostino-Pearson test, and comparisons were made using either unpaired Student’s t-test, or two-tailed Mann-Whitney U-test, as appropriate. Multiple comparisons were made using one-way ANOVA followed by Bonferroni’s post-hoc test. To analyze uniformity of distribution, the Hodjes-Ajne test for circular uniformity was used in MATLAB. P values were considered significant at P<0.05*, 0.01**, 0.001***, 0.0001****.

## RESULTS

### Effector T cells follow a Lévy-walk type of motility in the exocrine tissue

To study antigen-specific T cell behavior and motility patterns in the pancreas during autoimmune diabetes, we used the InsHA transgenic mouse model (34) in which fluorescent labels were introduced. We imaged influenza hemagglutinin (HA) antigen-specific CD8+ and CD4+ T cells attacking HA-expressing beta cells utilizing *in vivo* 2-photon microscopy (27,31). Co-transfer of naïve Clone 4-GFP CD8+ and HNT-CFP CD4+ T cells into sub-lethaly irradiated InsHA-mCherry hosts reproducibly induced pancreas infiltration by day 8 post-transfer (Fig. S1A), and hyperglycemia by day 10 (Fig. S1B). We were able to image beta cells, Clone 4-GFP CD8+ and HNT-CFP CD4+ T cells in pre-diabetic InsHA-mCherry mice and track T cell motility *in vivo* (Fig. 1A, Video S1). At day 8 post-transfer, HA-specific T cells in endocrine tissue (in islets) displayed lower average velocities than in the surrounding exocrine tissue (ref) and low directionality indexes (< 0.2) (ratio between cell’s displacement, defined as the straight line between original and final positions, and cell’s total track length) (Fig. 1B-C), as expected for T cells in presence of their cognate antigen (35). To describe T cell migration patterns, particle diffusion models have classically been used (32). T cells mostly migrate either following a Brownian-type random walk or a super-diffusive Lévy-type motility (characterized by stretches of directed motility in random directions interleaved by pauses) (4). Occasionally, T cells can display restrained motility (anomalous random walk or confinement) (8) or fully ballistic migration (in a straight path) (36), depending on imaging duration and the tissue analyzed. This models are based on the representation of cells mean square displacement (MSD) versus time (4). We fitted the experimental data with the different equations describing different models of diffusion (32) and identified the best fit based on the R^2^ values. While a complete Brownian-type random walk yields a linear regression between these parameters, a directed motility or a super-diffusive motility typical of a Lévy-walk are characterized by a power law curve, and confinement leading to sub-diffusive behavior yields a hyperbolic-shaped curve. Analyses of MSD of T cell populations versus time in islets revealed CD8+ T cell migration was best fitted with a model of confined motility, while CD4+ T cells migrated following a sub-diffusive (also called anomalous or restrained) random walk (Fig. 1D). In the exocrine tissue, mean T cell directionality index was in the 0.4 range (Fig. 1C), consistent with values reported for CTLs in a different model of insulitis (14), and indicative of an apparent lack of directionality. However, Clone 4-GFP and HNT-CFP T cell motility in the exocrine pancreas of InsHA-mCherry mice did not follow the described Brownian-type strictly diffusive random motility (14) and MSD of both T cell populations versus time (4) were best fitted with a model of super-diffusive Lévy-type motility, closely tending to a directed ballistic migration (36) (Fig. 1E-F, Video S2).

**Figure 1.**
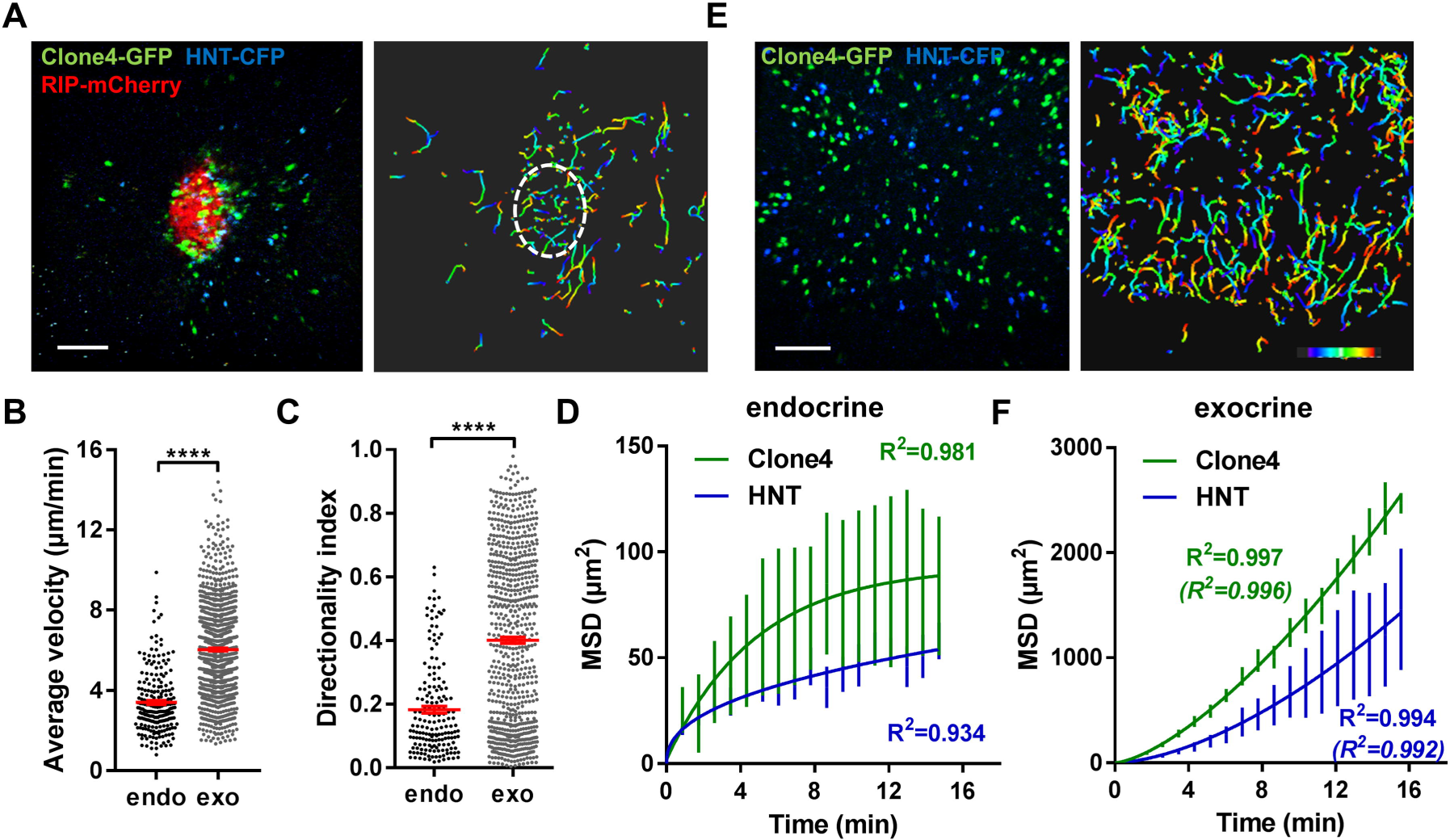
Motility of islet-antigen specific CD8+ and CD4+ T lymphocytes *in vivo*. Irradiated InsHA-mCherry mice adoptively transferred with Clone 4-GFP CD8+ and HNT-CFP CD4^+^ T cells were subjected to intra-vital microscopy on day 8. **A**) Still image from a representative movie (left panel; scale: 100 µm, 200 µm Z-projection; red: mCherry, green: GFP, blue: CFP) (See Video S1), and corresponding T cell tracks (right panel), color-coded as function of time. Islet is circled. Movie duration: 15 min. **B**) Average velocities of pooled CD4+ and CD8+ T cells in exocrine and endocrine tissues (n = 4 mice/condition; 1-2 movies/mouse, Mann-Whitney). Dots correspond to individual T cells. **C**) Directionality indexes (ratio between displacement and total track length) of T cells in exocrine and endocrine tissues (n = 4 mice/condition; 1-2 movies/mouse, Mann-Whitney). Dots correspond to individual T cells. **D**) Mean squared displacement (MSD) of T cells as function of time in islets, best fitted with a confined model of migration for Clone4-GFP cells, and with sub-diffusive random-walk for HNT-CFP cells. Bars correspond to SEM (n = 4 mice/condition; 1-2 movies/mouse). **E**) Still image from a representative movie in the exocrine tissue (left panel; scale: 100 µm, 200 µm Z-projection; green: GFP, blue: CFP) (See Video S2), and corresponding T cell tracks (right panel), color-coded as function of time. Movie duration: 19 min. **F**) MSD of T cells as function of time in the exocrine tissue, best fitted with a Lévy-walk model of migration. Between brackets are R^2^values of fit for ballistic (directed) motility. Bars correspond to SEM (n = 4 mice/condition; 1-2 movies/mouse).

### Contribution of chemotactic cues to T cell exploratory migration in the pancreas

Chemotaxis refers to the capacity of T cells to adapt their migratory pattern and motility following sensing of extrinsic cues produced by other immune cells or tissue specific cells. To analyze whether the super-diffusive motility in the exocrine tissue was informed by chemotactic cues produced within infiltrated islets, which are important sources of chemokines (37), and whether T cells were able to collectively migrate towards islets, we analyzed displacement of T cells towards (IN) or away (OUT) from islet centroids, as a function of T cell initial position (Fig. 2A-B). Proximity to islets did not bias T cell orientation of movement, as described in another model (14). Furthermore, although T cells migrated in rather straight paths in the exocrine tissue, cells did not collectively migrate in one particular direction in movies, as the distribution of compiled T cell vector orientations in different movies did not statistically differ from a uniform circular distribution (Fig. 2C-E). Thus, T cells do not seem to collectively answer to a large scale chemo-attractive gradient.

**Figure 2.**
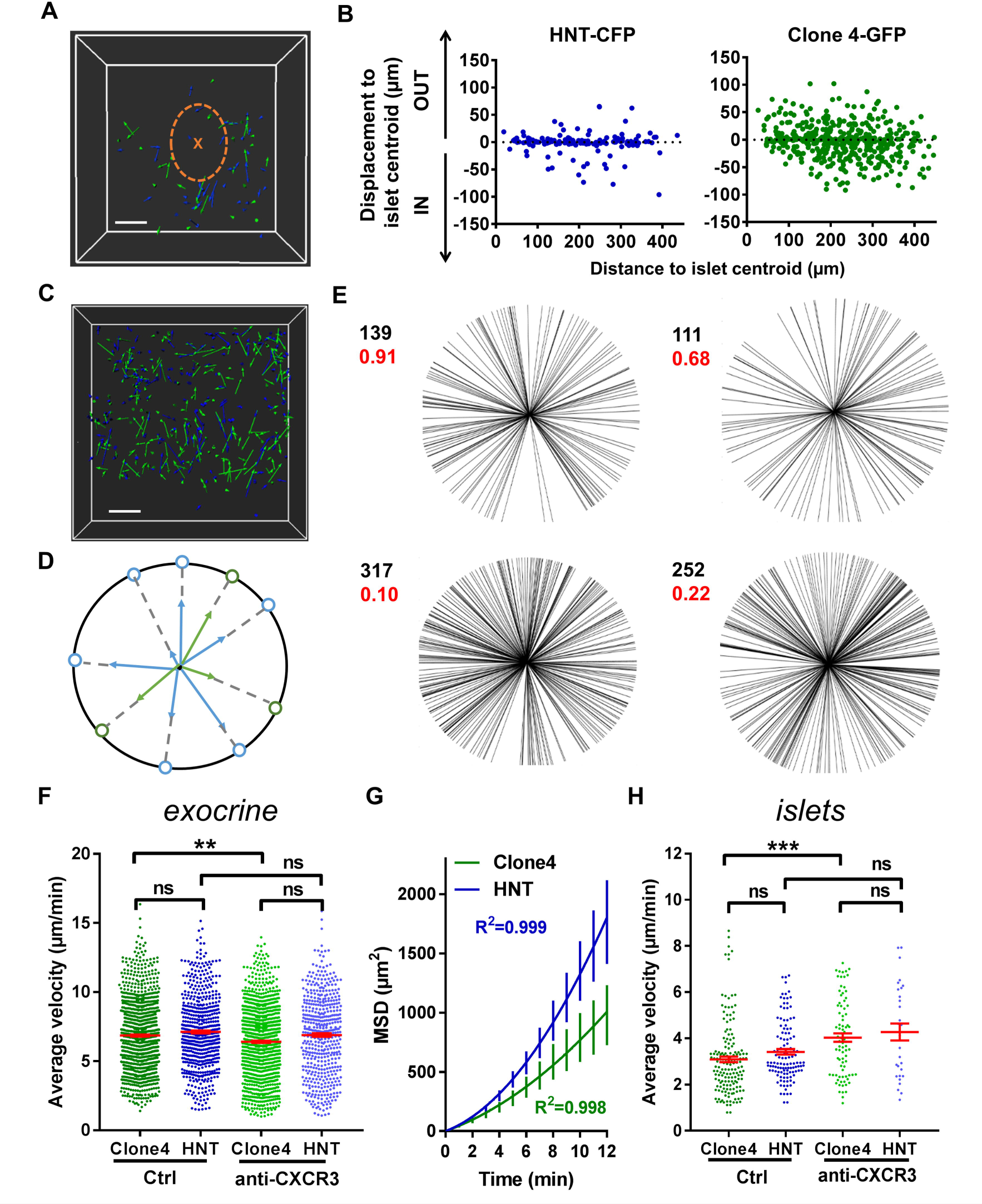
T cells collective migration is not biased towards islets and is mostly independent of CXCR3 signaling. Irradiated InsHA-mCherry mice adoptively transferred with Clone 4-GFP CD8+ and HNT-CFP CD4^+^ T cells were subjected to intra-vital microscopy on day 8. **A**) XY projections of track displacement vectors of T cells in movie in Fig. 1A (see Video S1). Scale: 100 µm. Blue and green tracks correspond to HNT-CFP and Clone 4-GFP T cells, respectively. Islet is circled, and a cross marks islet centroid. **B**) Clone 4-GFP and HNT-CFP displacement during movies towards (IN) or away (OUT) from islets, as function of distance from islet centroid at the start of movies (n = 4 mice; 1-2 movies/mouse). **C**) XY projections of T cell track displacement vectors in exocrine tissue (See Video S2) (scale: 100 µm, 200 µm Z-projection; green: GFP, blue: CFP). Movie duration: 10 min. **D**) To analyze orientations of T cell directions, displacement vectors were projected on the XY plane and set to a common origin. The orientation of each vector was projected on a circle. **E**) Statistical analysis of T cell track orientations in 4 different movies (without islet) (numbers in black correspond to number of tracks). None of the analyzed distributions were significantly different from a uniform distribution (Hodjes-Ajne test for circular uniformity, P values in red). **F**) Average velocities of CD4+ and CD8+ T cells in the exocrine tissue (n = 4-6 mice/condition; 2-3 movies/mouse, One-Way Anova). Dots correspond to individual T cells. CXCR3: anti-CXCR3 mAb-treated mice. **G**) MSD of T cells as function of time in the exocrine tissue, best fitted with a Lévy-walk super-diffusive model of migration. Bars correspond to SEM (n = 4 mice per condition; 1-2 movies/mouse). **H**) Average velocities of CD4+ and CD8+ T cells in islets (n = 3-6 mice/condition; 1 movie/mouse, One-Way Anova). Dots correspond to individual T cells.

An alternative possibility may be that T cells are able to respond to chemotactic cues locally (24). Indicative of this, T cells were able to follow each other for extended periods of time (> 5 min) in the exocrine tissue (Fig. S2A, Video S3). These events were detectable in all movies with infiltration > 100 cells/0.05 µm^3^ imaging volume (1-8 events/15 min movie, total 94 events over a total movie time of 4 h). Local cues may be either produced by the “leading” T cell, inducing other T cell to follow, or both T cells may be responding to the same chemo-attractive source. While T cells search for their cognate antigen, APCs are able to recruit them through secretion of different chemokines (38–40). CXCL10 is the most abundant chemokine expressed in infiltrated pancreas in mouse models, including InsHA, as well as in type 1 diabetic patients, and this chemokine contributes to T cell recruitment (37,41,42). The corresponding chemokine receptor CXCR3 was expressed by Clone 4-GFP CD8+ T cells infiltrating the pancreas and to a much lower extent by HNT-CFP CD4+ T cells (Fig. S2B). To determine whether signaling through this axis was involved in T cell migration in the pancreas, we treated transferred mice with anti-CXCR3 mAb 1h prior *in vivo* imaging. We found that this treatment had minor effects on T cell average velocities in the exocrine tissue (Fig. 2F) without changing the nature of migration statistics (Fig. 2G). In contrast, treatment with anti-CXCR3 mAb increased Clone 4-GFP CD8+ T cell motility in islets (Fig. 2H) and significantly reduced their recruitment into the pancreas (Fig. S2C). Thus, while CXCR3 has minor involvement in CD8+ and CD4+ T cell migratory pattern in the exocrine tissue it actively participates in CD8+ T cell recruitment and downregulates their velocity in islets, presumably to promote confinement of effector cells and local accumulation at sites of chemoattractant production (35) and/or cognate antigen presence.

### Blood vessels and ECM fibers provide a scaffold for T cell directed motility in the exocrine tissue *in vivo*

Diabetogenic T cells have been shown to extravasate and infiltrate the pancreas both within islets (24) and from post-capillary venules in the exocrine tissue (14). In accordance with this, early infiltration events here were limited to islets and perivascular areas (Fig. S3A). As infiltration progressed, large accumulations of effector CD4+ and CD8+ T cells could be observed within islets and at the level of endomucin-expressing pancreatic venules on fixed pancreas sections (Fig. S3B) and *in vivo* (Fig. S3C). Along large vessels (> 100 µm in diameter), T cells displayed linear tracks (Fig. 3A-D) (Video S4). To quantify alignment between T cell tracks and vessels, angle differences between track displacement vectors and vessel direction (1) (white lines, Fig. 3D) were measured for T cells close to or away from vessels (< 30 µm or > 30 µm) (Fig. 3E). Compared to other T cells in the imaging field, T cells in close proximity to vessels presented lower angle differences with vessels orientation, increased velocity, and fully ballistic motility (Fig. 3E-G). Thus, the vascular structure strongly influences all parameters of T cell migration in the pancreas.

**Figure 3.**
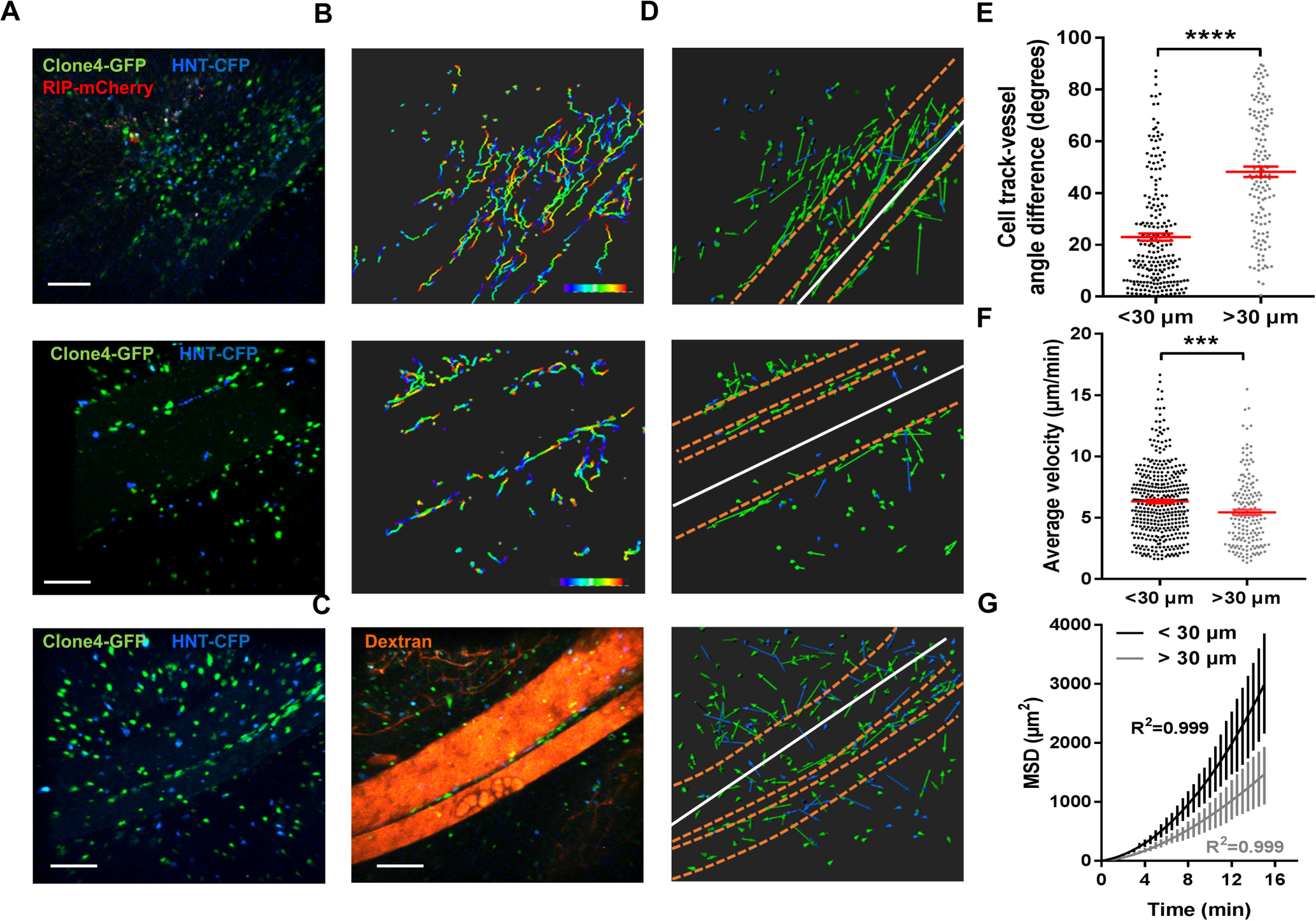
Blood vessels in the exocrine tissue contribute to effector T cells directed mode of motility. Irradiated InsHA-mCherry mice adoptively transferred with Clone 4-GFP CD8+ and HNT-CFP CD4^+^T cells were subjected to intra-vital microscopy on day 8. **A)** Still images from movies at day 8 post-transfer (Scale: 100 µm, 200-300 µm Z-projections) (See also Video S4). **B)** Corresponding T cell tracks in the imaging fields in (A), top and middle panels, color-coded as function of time. Movies duration: 19 min. **C)** Still image from the movie in (A) bottom panel post-injection of 150 kDa dextran-rhodamine (Scale: 100 µm, 300 µm Z-projection). **D)** XY projections of track displacement vectors of T cells in movies depicted in (A). Blue and green tracks correspond to HNT-CFP and Clone 4-GFP T cells, respectively. Dashed orange lines outline large vessels, and white lines indicate axis used to calculate angles between displacement vectors and vessel positions. **E)** Angle differences between displacement vectors of T cells close to vessels (pooled data from Videos S4, n = 3 movies from 3 mice) are lower than that of T cells away (> 30 µm) from vessels (Mann-Whitney). **F)** Average velocities of T cells close or away from vessels (> 30 µm) (n = 3 mice/condition; 1 movie/mouse, Mann-Whitney). Dots correspond to individual T cells. **G)** MSD of T cells close to (< 30 µm) vessels as function of time, best fitted with a directed model of migration, while other T cells follow a Lévy walk (> 30 µm from vessels). Bars correspond to SEM (n = 4 mice, 1 movie/mouse).

Different components of the ECM have been involved in guiding effector T cells migration through ligand-receptor interactions (1,15,16). Because vessels are usually lined with dense accumulation of ECM fibers, we analyzed T cell motility *in vivo* on ECM fibers visualized by second harmonic generation (SHG). We found the T cells were able to follow dense ECM bundles between vessels (Fig. 4A, Video S5). In addition, ECM fibers could be observed in the infiltrated exocrine tissue, although SHG was limited to the tissue surface (Fig. 4B, Video S6). Because ECM composition may be modulated by inflammation (25), we investigated whether pancreas infiltration was accompanied by changes in the ECM. Fibronectin, a key component of the ECM and major substrate for integrins (1), could be evidenced in the pancreas of non-treated control mice and localized to the perivascular space, as well as the interstitial tissue around cells in the exocrine pancreas (Fig. 4C). At day 8 post T cell transfer, we found an increase in fibronectin deposition at T cell infiltration sites (Fig. 4C). This was also true for other components of the ECM, such as collagen I (Fig. S4A). Importantly, assessment of Clone4-GFP and HNT-GFP T cells localization revealed a generalized close apposition to fibronectin fibers in pre-diabetic mice (Fig. 4D). Other changes in the micro-environment accompanying T cell infiltration, and locally correlated with fibronectin accumulation, included important APCs recruitment, as evidenced by dense CD11c and F4/80 labeling around and within islets and around blood vessels (Fig. S4B). Recruited T cells therefore migrate around a restructured scaffold of ECM fibers and leucocytes.

**Figure 4.**
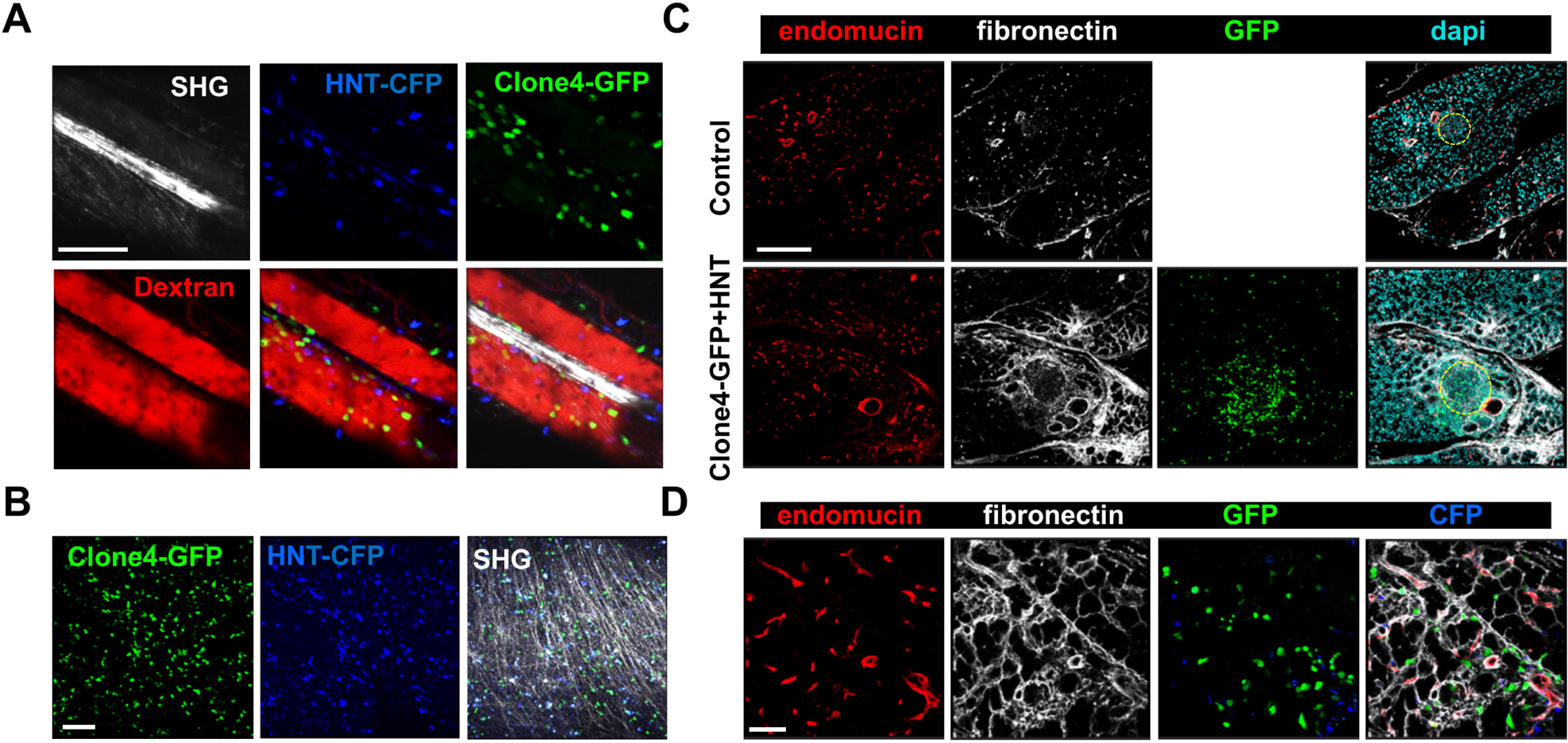
Effector T cells migrate along ECM fibers, which accumulate at infiltration sites. Irradiated InsHA-mCherry mice adoptively transferred with Clone 4-GFP and HNT-CFP T cells were subjected to intra-vital microscopy on day 8. **A)** Still images from a movie at day 8 post-transfer (Scale: 100 µm, 87 µm Z-projection) (See also Videos S5). SHG: second harmonic generation. Red: rhodamine-dextran. **B)** Still images from a movie at day 8 post-transfer (Scale: 100 µm, 100 µm Z-projection) (See also Videos S8). SHG: second harmonic. **C)** Representative confocal images of pancreas of a control irradiated InsHA-mCherry mouse, or post-transfer of T cells (scale: 200 µm, Z-projection of 20 µm). Islets are circled. **D)** Representative confocal images of exocrine tissue at day 8 post-transfer (scale: 50 µm, Z-projection of 4 µm), showing transferred T cells are in close apposition to fibronectin fibers.

### Integrin blockade alters directed effector T cell migration in the pancreas and impairs their effector phenotype

RGD binding integrins are known receptors for ECM proteins and in particular for fibronectin. Therefore, we assessed the expression of those that have been reported to be more frequently present on diabetogenic T cells (1) in the infiltrating effector T cells of our model. We found that the vast majority of both Clone 4-GFP CD8+ and HNT CD4+ T cells expressed high levels of β_1_ and αV integrins (Fig. S5). We hypothesized that integrins could be involved in guiding effector T cell motility in the pancreas. We tested this hypothesis by injecting a blocking anti-β_1_ integrin mAb and found that shortly post-injection (40 min), average velocities of both Clone 4 and HNT T cells were significantly reduced compared to isotype control antibody-treated animals (*20 %) (Fig. 5A-C, Video S7), as well as the directionality indexes of T cell tracks (Fig. 5D). In addition, although T cells MSD versus time curves were still best fitted with the Lévy-type random model, curves tended to linearize and the fit for a Brownian-type random motility improved in treated animals (Fig. 5E). To further assess involvement of integrins in T cells motility in the pancreas, we treated animals prior to imaging (10 min) with a peptide containing the RGD peptidic motif. Since this sequence is recognized by integrins on ECM fibers, treatment with RGD peptide broadly blocks integrins. We found that average T cell velocity was decreased compared to reverse DGR peptide treated-animals (Fig. S6A-B, Video S8) and super-diffusive motility was practically lost (Fig. S6C). This indicates that integrins contributes to T cell motility in the inflamed pancreas, although compensatory and/or additional mechanisms may exist (17,20).

**Figure 5.**
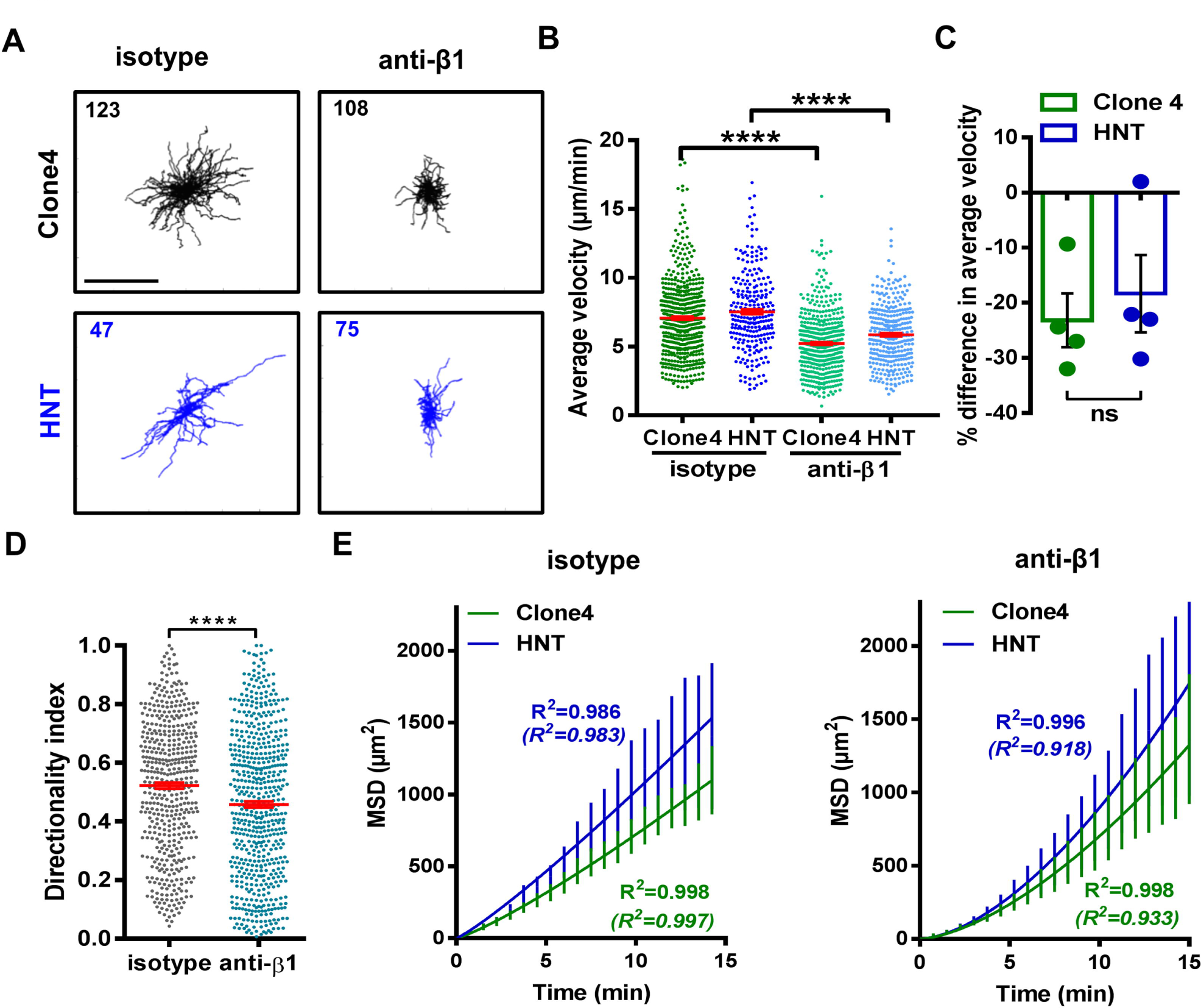
β_1_ integrin-dependent interactions between T cells and the ECM shape T cell motility. Irradiated InsHA-mCherry mice transferred with Clone 4-GFP and HNT-CFP T cells were subjected to intra-vital microscopy on day 8. **A**) XY projections of T cell tracks over 15.2 minutes 35 min after control IgG or anti-β_1_ integrin injection (scale: 100 µm) (See also Video S7). Values indicate number of tracks in movies. **B**) Average velocity of T cells in the exocrine tissue of isotype and anti-β_1_ integrin treated animals (n = 3 mice/condition; 1-2 movies/mouse; Mann-Whitney). Dots correspond to individual cells. **C**) Percentage difference in average velocity between T cells in the exocrine tissue of isotype and anti-β_1_ integrin-treated animals. Dots correspond to individual movies (n = 3 mice/condition; 1-2 movies/mouse; P = 0.48, Mann-Whitney). **D)** Directionality index of T cells in exocrine tissue (n = 3 mice/condition; 1-2 movies/mouse, Mann-Whitney). **E)** MSD of T cells as function time in the exocrine tissue of isotype and anti-β_1_ integrin-treated animals were both best fitted with a model of Lévy-type super-diffusive migration (solid lines). Between brackets are R^2^ values of fit for Brownian random motility. Bars correspond to SEM (n = 3 mice/condition; 1-2 movies/mouse).

Finally, we tested whether impaired effector T cell motility induced by β_1_ integrin blockade could affect functionality. Mice were treated with anti-β_1_ integrin mAb at a time at which T cells had already infiltrated the pancreas (days 8 and 9 after transfer). At day 10, equal numbers of infiltrating Clone 4-GFP CD8+ and HNT-CFP CD4+ T cells were detected in the pancreas of treated compared to isotype control mice (Fig. 6A). Moreover, the phenotype and cytokine secretion potential of both donor CD8+ and CD4+ T cells were indistinguishable in the draining lymph nodes of the pancreas (Fig 6B-C). These results indicate that treatment at this time point did not prevent activation and recruitment of effector cells into the pancreas. In contrast, pancreas infiltrating HNT-CFP CD4+ T cells from treated mice displayed a significant reduction in the expression of key effector markers such as KLRG1 and CD25 (Fig. 6B). Additionally, these cells had lost the potential to secrete IL-2, an important effector cytokine (Fig. 6C). Interestingly, although the effector potential of Clone 4-GFP CD8+ T cells remained unaltered, a marked reduction of the expression of CD25 was observed in treated mice, likely as a result of the decreased IL-2 secretion by helper CD4+ T cells (Fig. 6C). Collectively, our data indicate that altered motility of diabetogenic T cells in the pancreas results in deceased effector functions *in situ*.

**Figure 6.**
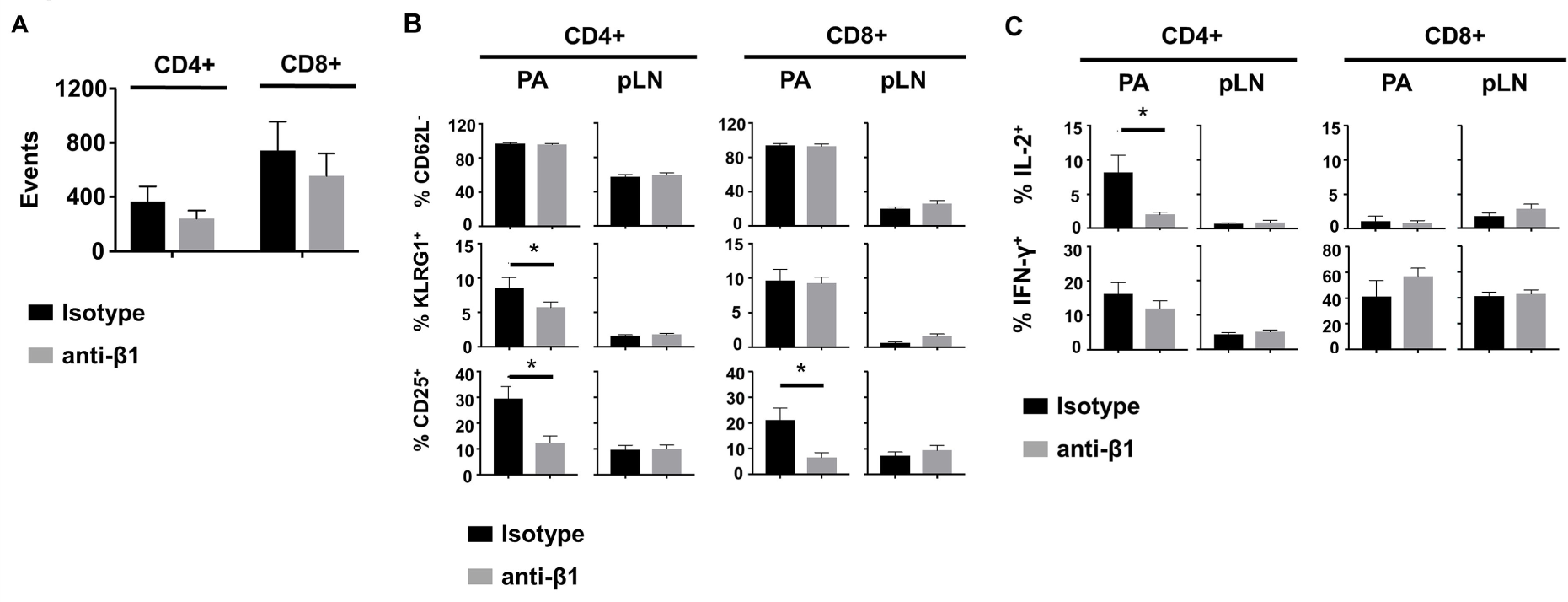
β_1_ integrin blockade alters diabetogenic T cell effector phenotype in the pancreas. Irradiated InsHA-mCherry mice transferred with Clone 4-GFP CD8+ and HNT-CFP CD4+ T cells were treated with anti-β1 mAb or isotype control antibodies on days 8 and 9 after transfer. At day 10, donor T cells from pancreatic LN (pLN) and pancreas (PA) were analyzed by FACS gating on living CD8+ or CD4+ Thy1.1+ lymphocytes. **A**) Donor T cells in the pancreas of treated mice. FACS event counts in the CD8+ or CD4+ Thy1.1+ gates from 3 independent experiments, represented as mean ± SEM (n = 8 mice, Mann-Whitney). **B**) Donor T cells expression of CD25, CD62L and KLRG1. Percentages of indicated subpopulations in the CD8+ Thy1.1+ or CD4+ Thy1.1+ gates from 3 independent experiments, represented as mean ± SEM (n = 8 mice, Mann-Whitney). **C**) Intracellular cytokine measurement in donor T cells. Percentages of IL-2+ or IFNΓ+ cells in the CD8+ Thy1.1+ or CD4+ Thy1.1+ gates from 2 independent experiments, represented as mean ± SEM (n = 6 mice, Mann-Whitney).

## DISCUSSION

T cell migratory behavior stems from the need to search for their cognate antigen and plays a crucial role in antigen clearance. In various peripheral tissues (brain, liver, gut, pancreas), T cell migratory behavior has been described as a super-diffusive random walk or Lévy walk (4,14,40), characterized by steps of directed migration in random directions interleaved by pauses (43), to optimize rare target encounter. However, mechanisms governing T cell migration are context-dependent and leukocytes are able to switch migratory modes along with changing environmental conditions (1). This prevents the definition a generalized model for T cell interstitial migration in inflamed peripheral tissues. In addition, mechanisms governing lymphocyte dynamics are intimately linked to the maintenance of T cell effector function (1). While effector T cells need to reach dispersed target islets in the pancreas during autoimmune diabetes, mechanisms governing their motility remained unclear. Using 2-photon microscopy *in vivo* to visualize TCR transgenic HA-specific CD8^+^ and CD4^+^ T cells in the pancreas of mice expressing HA in beta cells, we found that both T cell types followed a super-diffusive Lévy-type mode of migration in the exocrine tissue without a preferred concerted orientation. In contrast, the islet environment restrained T cell trafficking through a mechanism involving CXCR3 chemokine receptor. T cell infiltration induced local fibrosis, marked by fibronectin deposition. Both CD8^+^ and CD4+ T cells were in close apposition to vessels and fibronectin fibers, which provided adhesive guidance and contributed to the super-diffusive migration in the exocrine pancreas through, at least partially, an integrin-dependent mechanism. Finally, integrin-dependent T cell-ECM interactions contributed to the maintenance of T cell effector function in the pancreas.

In search for their cognate antigen, T cells can follow a “Brownian” random walk mode of migration in the periphery, including in the pancreas (6,14). Here, however, T cells followed a super-diffusive, almost ballistic, mode of migration (4), which arguably constitutes the most efficient strategy of random search processes (44). The different migration pattern observed in the exocrine tissue by Coppieters et al. may rise from model-specific differences (autoimmune diabetes used was induced using viral infection) and/or different length of movie duration to analyze T cell MSD (< 7 min vs. 12-16 min here) (14). Strikingly, T cells did not collectively migrate in a particular direction as no overall orientation bias of T cell tracks was observed, including towards islets, although these are major sources of chemokines (37). We found that, unlike in LN (45), Gαi-coupled receptors involved in chemokine signaling, such as CXCR3 receptors, were not central in shaping T cell motility in the exocrine pancreas. CXCR3 blockade slightly reduced T cell velocity without affecting migration mode, as reported previously in the brain (43). On the local scale, the fact that T cells were able to follow each other suggests that they may follow paths of least resistance. Alternatively, like recently described for neutrophils (46), T cells may be able to deposit chemokine trails that other T cells may be able to respond to, although this remains unclear.

Although the original assumption was that large scale diffusive chemokine gradients would provide cues for directed motility, experimental evidence of collective T cell migration towards sources of high chemokine production is scarce. In contrast, chemotactic cues are able to modulate T cell trajectories in different ways, such as through modulation of T cell retention/arrest rather than directionality (35). In accordance with this, large accumulation of T cells were observed in islets and CXCR3 blockade increased T cell velocity in islets, as beta cells are the main source of CXCL9/10 in the pancreas (37) The chemokine-rich environment of islets therefore promotes a downregulation of T cell velocity to accumulate and confine effector cells at target sites, rather than attract distant T cells. The dense accumulation of T cells at the level of post-capillary venules in the exocrine tissue could be explained by the described vascular leakiness (14), and presence along vessels of CD11c+ and F4/80+ cells, which are well-known chemokine sources that could favor confinement of T cells.

Similar to what was described in the inflamed skin (1), the vascular tree provided a scaffold for T cell migration in the pancreas and strongly contributed to the directional motility *in vivo*. In addition, HA-specific CD8+ and CD4+ T cell infiltration induced ECM remodeling, likely mediated by recruited macrophages (47). This remodeling included fibronectin accumulation, a major substrate for integrins (1). The fact that anti-β1 integrin mAb treatment affected both velocity and directionality of T cells indicates that lymphocytes do not only align along paths of least resistance in the pancreas, but that fibronectin fibers also provide adhesive guidance. Effects observed were in line with previous studies of integrin-blockade on T cell motility (48). By contrast with full integrin-dependency described in the inflamed skin (1), our results suggest the implication of complementary mechanisms of migration for T cells in the pancreas. Once the described chemokine-dependent up-regulation of integrin molecules allowing T cell entry at peripheral sites has been achieved (49), infiltrated T cell directional migration in the pancreas is mostly independent of CXCR3-mediated chemotactic signals. Remaining migration detected in the presence of integrin-blocking antibody may stem from T cells intrinsic capacity to maintain a directed motion (50), other GPCR-mediated chemokine signaling (51), and/or other receptor-ligand interactions, although this remains to be clarified.

Finally, integrin β1 blockade at a time when diabetogenic T cells had already infiltrated the pancreas resulted in a decline of CD4+ T cell effector function. Diabetogenic T cells require antigen-mediated contacts with APCs in the pancreas to retain effector function over time (27,52). A likely explanation may be that impairing ECM guided motility would alter effector CD4+ T cell/APC interactions resulting in decrease in effector function. Alternatively, integrin signaling triggered by direct interaction with ECM fibers might be required for the maintenance of effector functions in the pancreas, as described in other settings (53,54).

In summary, we show that during autoimmune insult to the pancreas, islet-antigen specific T cells display super-diffusive motility in the exocrine tissue implicating integrin-dependent T cell-ECM fibers interactions contributing to optimization of islet encounter and maintenance of effector functions, and that the islet chemokine-rich environment promotes the confinement of effector T cells, rather than their recruitment. We thus reveal a role for integrins in the pancreas that may have important implications for the design of new therapeutic strategies against T1D.

## AUTHOR CONTRIBUTIONS

GEC, CLS, JH and MS designed experiments; GEC, CLS, AM and MS performed experiments; GEC, CLS, PF, JH, and MS analyzed data; PM, JH and MS wrote the manuscript.

## ACKNOWLEDGMENTS

The authors declare no conflict of interest. The authors would like to thank C. Lafont, P. Samper and E. Galibert, Institute of Functional Genomics, Montpellier, France, for technical assistance, the animal facility staff (RAM), and Montpellier Imaging Platform (IPAM). Authors were supported by grants from the Agence Nationale de la Recherche (ANR BETA-DYN JCJC13 to MS, ANR MITOSTEM to JH, France-BioImaging ANR-10-INBS-04 to PM), Société Francophone du Diabète, INSERM, CNRS, University of Montpellier, and Région Occitanie.

